# Photoreceptor genes in a trechine beetle, *Trechiama kuznetsovi*, living in the upper hypogean zone

**DOI:** 10.1101/2023.02.28.530396

**Authors:** Takuma Niida, Yuto Terashima, Hitoshi Aonuma, Shigeyuki Koshikawa

## Abstract

To address how organisms adapt to a new environment, subterranean organisms whose ancestors colonized subterranean habitats from surface habitats have been studied. Photoreception abilities have been shown to have degenerated in organisms living in caves and calcrete aquifers. Meanwhile, the organisms living in a shallow subterranean environment, which are inferred to reflect an intermediate stage in an evolutionary pathway to colonization of a deeper subterranean environment, have not been studied well. In the present study, we examined the photoreception ability in a trechine beetle, *Trechiama kuznetsovi*, which inhabits the upper hypogean zone and has a vestigial compound eye. By *de novo* assembly of genome and transcript sequences, we were able to identify photoreceptor genes and phototransduction genes. Specifically, we focused on *opsin* genes, where one *long wavelength opsin* gene and one *ultraviolet opsin* gene were identified. The encoded amino acid sequences had neither a premature stop codon nor a frameshift mutation, and appeared to be subject to purifying selection. Subsequently, we examined the internal structure of the compound eye and nerve tissue in the adult head, and found potential photoreceptor cells in the compound eye and nerve bundle connected to the brain. The present findings suggest that *T. kuznetsovi* has retained the ability of photoreception. This species represents a transitional stage of vision, in which the compound eye regresses, but it may retains the ability of photoreception using the vestigial eye.

## Background

How organisms adapt to a new environment is one of the fundamental research questions in evolutionary biology [1]. Subterranean organisms whose ancestors originally lived in a surface environment are ideal for investigating this issue [2, 3]. Subterranean habitats are not continuously exposed to light, and can be categorized into cave habitats, interstitial habitats and superficial subterranean habitats [4, 5]. Degeneration of eyes is generally observed in various taxa colonizing these subterranean environments [6, 7].

Do the organisms having a regressed eye also have decreased ability of photoreception? Previous studies focused on various aspects of subterranean adaptation, including signatures in photoreceptor proteins [8, 9]. For example, one previous study described the expression of visual opsin genes which encode seven-transmembrane photoreceptor proteins in the binocular eye of the Mexican blind cavefish, *Astyanax mexicanus* [8]. Transcripts of visual opsin genes were underrepresented in the cavefish as compared with a conspecific surface population, and this could be attributed to reduction of photoreceptor cells in the cavefish [10].

In Insecta, subterranean diving beetles (Dytiscidae), which have highly regressed or no eyes and inhabit a calcrete aquifer located 10 m underground in Western Australia, were subjected to a similar analysis [11]. Transcripts were not detected for *long wavelength opsin* or *ultraviolet opsin* at the adult stage of the diving beetles [12], and pseudogenization of *long wavelength opsin, ultraviolet opsin* and some phototransduction genes (i.e. *arrestin, neither inactivation nor afterpotential C, transient receptor potential* and *transient receptor potential channel-like*) was observed [13, 14].

Besides calcrete aquifers, insects have also been found to colonize superficial subterranean habitats, such as rock fissures near the surface [15]. Insects living in a superficial subterranean habitat can be exposed to light due to unexpected environmental fluctuation and migration of insects themselves toward the surface, unlike insects living in deep subterranean habitats. The colonization of a superficial subterranean habitat is inferred to reflect an intermediate stage in an evolutionary pathway to colonization of deeper and extreme environments [4]. Despite this importance, the biological features of species living in a superficial subterranean habitat remain unexplored.

The present study focused on one trechine beetle species, *Trechiama kuznetsovi* (Coleoptera: Carabidae: Trechinae). This species has a vestigial compound eye and inhabits the upper hypogean zone, which is a similar environment to mesovoid shallow substratum (MSS) [16]. We aimed to reveal the photoreception ability in this species. We obtained genome and transcript sequences to examine photoreceptor genes and phototransduction genes and estimate selective pressure on visual opsin genes. Also, histological investigations were performed to observe the internal structure of the vestigial compound eye and a nerve bundle connecting it to the brain.

## Methods

### Sample collection

*Trechiama kuznetsovi* samples were collected at Yûbari City, Hokkaido, Japan. We collected *T*. *kuznetsovi* adults from the upper hypogean zone consisting of small rocks and clay, by digging in the slope of a v-shaped valley to the depth of a few to some dozen centimeters (Fig. 1).

**Fig. 1.**
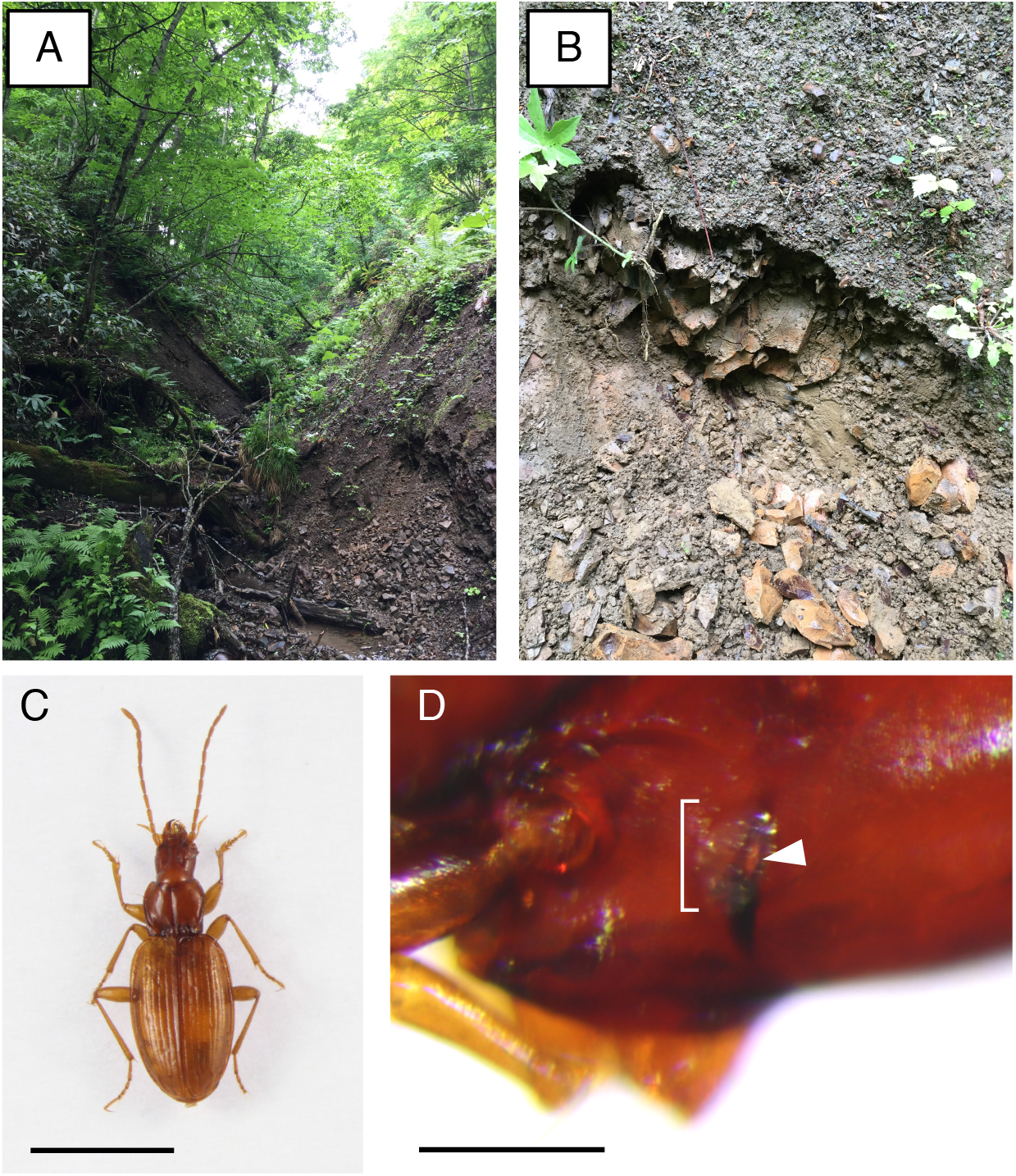
The habitat and morphology of *T. kuznetsovi*. (A) A v-shaped valley of a mountain area at Yûbari City in Hokkaido. (B) The upper hypogean zone under the slope of a v-shaped valley. It consists of small rocks and clay. (C) Dorsal view of *T. kuznetsovi*. Scale bar indicates 3 mm. (D) Lateral view of *T. kuznetsovi* head. The bracket indicates the vestigial compound eye. The black part is the pigmentation of the cuticle behind a compound eye (ocular ridge). The arrowhead indicates a hole on the ocular ridge, through which the nerve bundle is presumed to pass. There is no pigmentation in cells inside the compound eye structure. Scale bar indicates 0.2 mm.

### DNA and RNA sequencing

We used one adult male of *T. kuznetsovi* stored in 99.5% ethanol for genome sequencing. Before DNA extraction, mites adhering to the body surface were removed and the male genitalia was preserved in 99.5% ethanol for identification. Genomic DNA was extracted using a Wizard Genomic DNA Purification Kit (Promega, Madison, WI, USA). A library was constructed using a TruSeq Nano DNA Library Prep Kit (Illumina, San Diego, CA, USA) and sequenced on the NovaSeq 6000 platform (Illumina) by Macrogen Service (Macrogen, Seoul, South Korea). 2 × 151 bp paired-end reads were generated (Table S1).

We used one live adult male of *T*. *kuznetsovi* for transcript sequencing. Before RNA extraction, the beetle was washed with 99.5% ethanol, mites adhering on its body surface were removed and the male genitalia was preserved in 99.5% ethanol for identification. Total RNA was immediately extracted using an RNeasy Micro Kit (Qiagen, Hilden, Germany). A library was constructed using a SMARTer Stranded RNA-Seq Kit (Illumina) and sequenced on the NovaSeq 6000 platform (Illumina) by Macrogen Service. 2 × 101 bp paired-end reads were generated (Table S1).

### Assembly and mapping

Summary statistics of raw reads and adapter contamination were checked using FastQC (v0.11.9; Babraham Institute). Quality control was performed using fastp v0.20.1 [17] and Trimmomatic v0.39 [18] to trim off one base from the 3’ end, low quality sequences and adapter sequences. Then, summary statistics were rechecked using FastQC. The kmer content of reads from genome sequencing and the genome size were calculated using KmerGenie v1.7051 [19]. For the reads that passed the quality control, genome and transcript assembly were conducted with Platanus v1.2.4 [20] and Trinity v2.8.4 [21, 22]. In the scaffolding step of genome assembly, contigs smaller than 500 bp were excluded [23]. Completeness of the assembled genome and transcript was assessed using BUSCO_v5 for insecta core gene sets and CEGMA for invertebrate core gene sets in gVolante web server [24]. Summary statistics of the assembled sequences were calculated using SeqKit v.0.16.1 [25].

RNA reads were mapped to the assembled genome scaffolds using HISAT v2.1.0 [26]. The result of the mapping was visualized using IGV v2.11.9 [27].

### Homology search

We conducted a search for genes encoding three photoreceptor proteins and 15 phototransduction proteins from the assembled genome and transcripts with tblastn program using the following queries (unless noted, protein sequences of *Tribolium castaneum* [Tenebrionidae] were used): Long wavelength opsin (Lw opsin) of *Pogonus chalceus* (Carabidae: Trechinae), Ultraviolet opsin (Uv opsin) of *P. chalceus*, C-opsin, Arrestin 1 (Arr1), Arrestin 2 (Arr2), G protein α-subunit 49B (Gα49B), G protein β-subunit 76C (Gβ76C), G protein γ-subunit 30A (Gγ30A), G protein-coupled receptor kinase 1 (Gprk1), Inactivation no afterpotential D (InaD), Neither inactivation nor afterpotential A (NinaA), Neither inactivation nor afterpotential C (NinaC), No receptor potential A (NorpA), Protein C kinase 53E (Pkc53E) of *Drosophila melanogaster*, Rab-protein 6 (Rab6), Retinal degeneration C (RdgC), Transient receptor potential (Trp), and Trp-like (Trpl). Selection of these proteins was based on previous studies [28, 29]. Blast-hit sequences with an e-value < 1 × e^-20^ were treated as having high similarity [30]. If no sequence matched this criterion, Blast-hit sequences were examined in order from the best-hit sequence. In BLAST-search for the transcripts, the presence of premature stop codons and frameshift mutations was examined.

We subsequently conducted a further analysis of *lw opsin* and *uv opsin*, which are visual photoreceptor genes in Coleoptera, while some opsin genes are known to have light-independent roles in *D. melanogaster* [31–35]. We checked whether the blast-hit transcripts matched exon regions with mapped short reads of transcripts by HISAT. Matched transcripts were used for subsequent comparative analyses with a related surface species: *P. chalceus*, whose opsin amino acid sequences were already registered in NCBI protein database [32].

### Identification of *lw opsin* gene

A part of the genome sequence that has high similarity score to the Lw opsin amino acid sequence of *P. chalceus* was divided into three scaffolds (Table S2). To join these scaffolds together, primers were designed on each scaffold using Primer3Plus (https://www.bioinformatics.nl/cgi-bin/primer3plus/primer3plus.cgi) (Table S3) and PCR was performed using PrimeSTAR Max DNA Polymerase (Takara Bio, Shiga, Japan). The sequences of the PCR products, which were determined by Sanger sequencing, were overlapped on the scaffolds. The transcript sequence that showed similarity to the Lw opsin amino acid sequence of *P. chalceus* using tblastn was divided into two contigs (Table S2). These contigs were joined together by Sanger sequencing in the same method as above. Primers were designed on each contig (Table S3) and RT-PCR was performed using a 3’ RACE CORE Set (Takara Bio).

To specify exon and intron regions, the acquired transcript sequence of the *lw opsin* gene was aligned to the acquired genome sequence of the *lw opsin* gene with Exonerate v2.4.0 [36]. The exon and intron regions were illustrated with GenePalette [37].

### Identification of *uv opsin* gene

There was one genome scaffold that had high similarity score to the Uv opsin amino acid sequence of *P. chalceus* using tblastn (Table S2), but no transcript contig was found. However, short read sequences originated from RNA were mapped to the scaffold of the *uv opsin* gene. We determined the transcript sequence of the *uv opsin* gene by Sanger sequencing. Firstly, the exon regions were predicted with Exonerate using the Uv opsin amino acid sequence of *P. chalceus*. Then primers were designed on the predicted exon regions with Primer3Plus (Table S3) and RT-PCR was performed using a 3’ RACE CORE Set.

To specify exon and intron regions, the acquired transcript sequence of the *uv opsin* gene was aligned to the acquired genome sequence of the *uv opsin* gene with Exonerate. The exon and intron regions were illustrated with GenePalette.

### Opsin phylogeny and tests of selection

Blastp was performed using the acquired amino acid sequences of opsins in *T. kuznetsovi* as queries for non-redundant protein sequences in the NCBI database (Table S4). Amino acid sequences of opsins in *T. kuznetsovi*, four beetle species (*P. chalceus, Gyrinus marinus, Thermonectus marmoratus* and *T. castaneum*) and a honeybee (*Apis mellifera*) were aligned with MUSCLE in MEGA v 11 [38]. Based on the maximum likelihood method, a phylogenetic tree of nucleotide sequences was reconstructed under the best-fit GTR + G + I model with 1000 bootstrap generations.

The ancestral sequences of opsin gene sequences between subterranean *T. kuznetsovi* and surface *P. chalceus* were estimated, based on above five beetle’s phylogenetic relationship using MEGA. Based on the maximum likelihood method [39], the ratios of nonsynonymous (Ka) to synonymous (Ks) nucleotide substitution rates were calculated between an ancestral sequence and a sequence of *T. kuznetsovi* and between the ancestor sequence and a sequence of *P. chalceus* using KaKs_Calculator v 3.0 [40]. Fisher’s exact test on a 2 × 2 contingency table was conducted using the number of synonymous and nonsynonymous sites and synonymous and nonsynonymous substitutions.

The Ka/Ks analysis is able to suggest that observed changes in a sequence have been influenced by positive selection (Ka/Ks > 1), neutral evolution (Ka/Ks = 1), or negative (purifying) selection (Ka/Ks < 1). In our study, the apparent result is expected to be that opsin genes of *T. kuznetsovi* were under purifying selection, because along the evolutionary branch from an ancestor to *T. kuznetsovi*, opsins will have been selected under surface habitat before this lineage colonized subterranean habitat. To resolve this problem, we also compared the degrees of purifying selection between opsin genes of *T. kuznetsovi* (test) and *P. chalceus* (reference), carrying out branch-by-branch analyses with RELAX in Hyphy [41]. As the result of this model, *k* < 1 is indicative of relaxed selection, while *k* > 1 is indicative of purifying selection.

### Histological study

The internal structure of a vestigial compound eye in *T. kuznetsovi* adults was observed using paraffin sections. Adult heads were fixed in 50% alcohol Bouin solution (ethanol:Bouin solution [for pathology, Fujifilm Wako Pure Chemicals] = 1:1) at room temperature overnight or longer. The fixed samples were rinsed in 70% ethanol, dehydrated in increasing concentrations of ethanol (90%, 95% and 100%) [42, 43], and then cleared in xylene. Next, the samples were embedded in paraffin (Paraplast Plus; Sigma Aldrich, MO, USA), and transverse sections (6 μm) were serially cut with a microtome (OSK 97LF506; Ogawa Seiki, Tokyo, Japan). Sections were stained with hematoxylin, observed with a microscope (CX-43; OLYMPUS, Tokyo, Japan) and photographed with a mounted camera (EOS Kiss X9; Canon, Tokyo, Japan).

Because we could not obtain the complete series of cross sections due to their friability, the nerve bundle between a compound eye and a brain in *T. kuznetsovi* adults was observed with dissection. Adult heads were dissected in phosphate-buffered saline and stained with 0.5 % methylene blue solution (22409-32; NAKALAI TESQUE, Kyoto, Japan) for 1 hour. The samples were washed in phosphate-buffered saline, observed with a stereo microscope (SZX16; OLYMPUS) and photographed with a mounted camera (EOS Kiss X9; Canon).

## Results

### DNA and RNA sequencing

The genome size was estimated to be 554,652,206 bp based on the k-mer frequency distribution of genome reads with KmerGenie. The assembled genome contained 55,616 scaffolds with a total length of 456,726,283 bp (N50: 16,592 bp) and 96.20 % BUSCO completeness (Table 1). The assembled transcripts contained 71,303 contigs with a total length of 67,456,903 bp (N50: 1,883 bp) and 89.10 % BUSCO completeness (Table 1). 81.90% of RNA reads were mapped to the assembled genome with HISAT. The information of paired-end reads is summarized in Table S1.

**Table 1.** The summary statistics for assembly of the genome and transcripts in *T. kuznetsovi*.

|  | Genome | Transcripts |
| --- | --- | --- |
| Total assembled length (bp) | 456,726,283 | 67,456,903 |
| Scaffolds (n) | 55,616 | 71,303 |
| The longest scaffold (bp) | 306,214 | 52,805 |
| The shortest scaffold (bp) | 386 | 181 |
| Scaffold N50 (bp) | 16,592 | 1,883 |
| BUSCO pipeline analysis<br>(% complete / partial) | 96.20 / 98.83 | 89.10 / 95.32 |
| CEGMA pipeline analysis<br>(% complete / partial) | 93.55 / 100 | Not Available |

### Expression of photoreceptor genes and phototransduction genes

In the assembled genome, BLAST search found *lw opsin* gene, *uv opsin* gene and non-visual *c-opsin* gene (Table 2). *lw opsin* and *c-opsin* were found in the assembled transcripts, but *uv opsin* was not. Transcripts for 14 of the 15 BLAST-searched phototransduction genes were detected, namely, *Arr1, Arr2, Gα49B, Gβ76C, Gγ30A, Gprk1, inaD, ninaA, ninaC, norpA, Pkc53E, Rab6, rdgC* and *trp*. Most of these transcript sequences had neither a premature stop codon nor a frameshift mutation, but transcript sequences of *Gα49B, Gβ76C, Gprk1, norpA* and *rdgC* had premature stop codons or frameshift mutations in some transcript isoforms. These transcripts might include primary transcripts before splicing. One phototransduction gene, *trpl* was not detected in the transcripts, but was found in the genome.

**Table 2.**
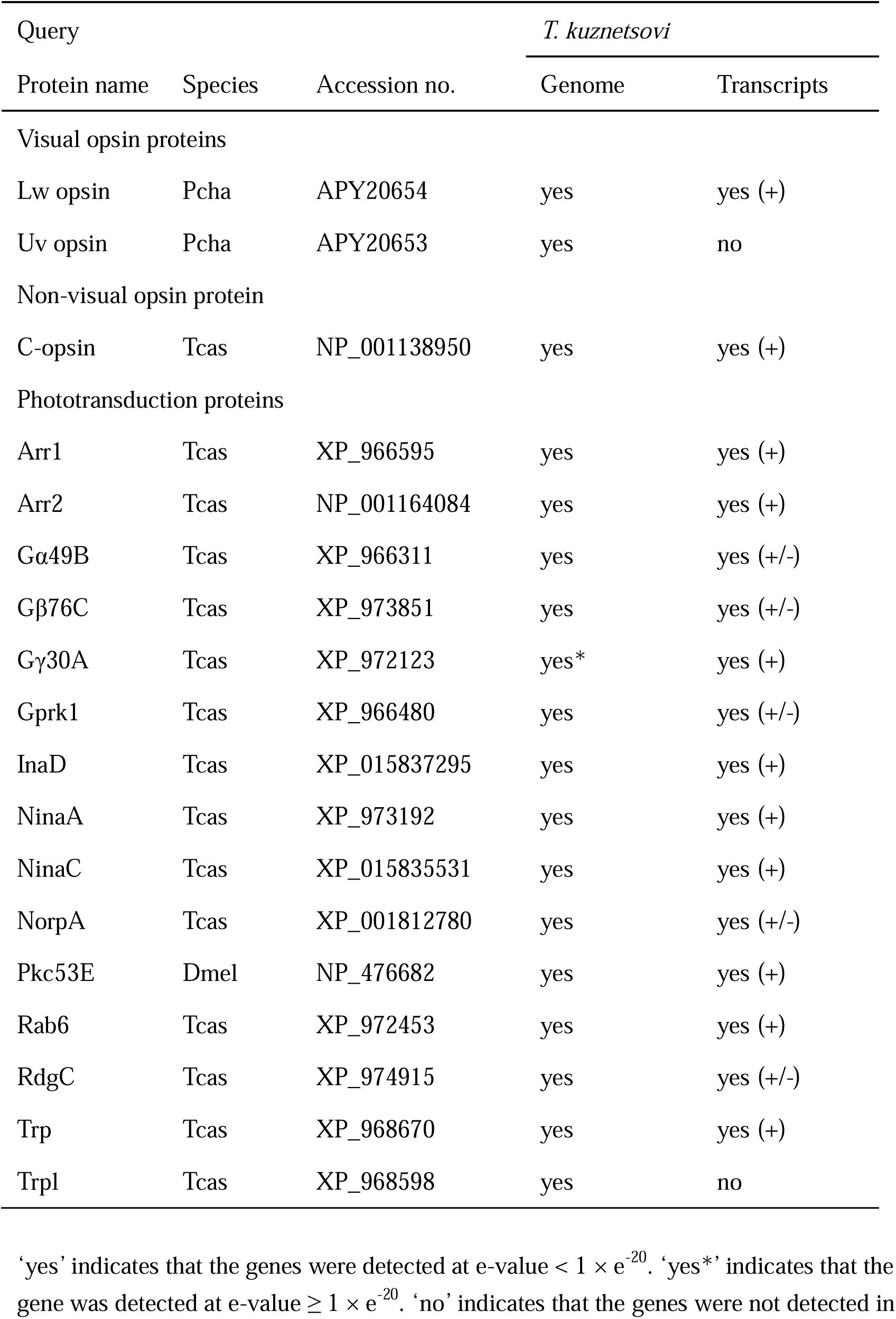
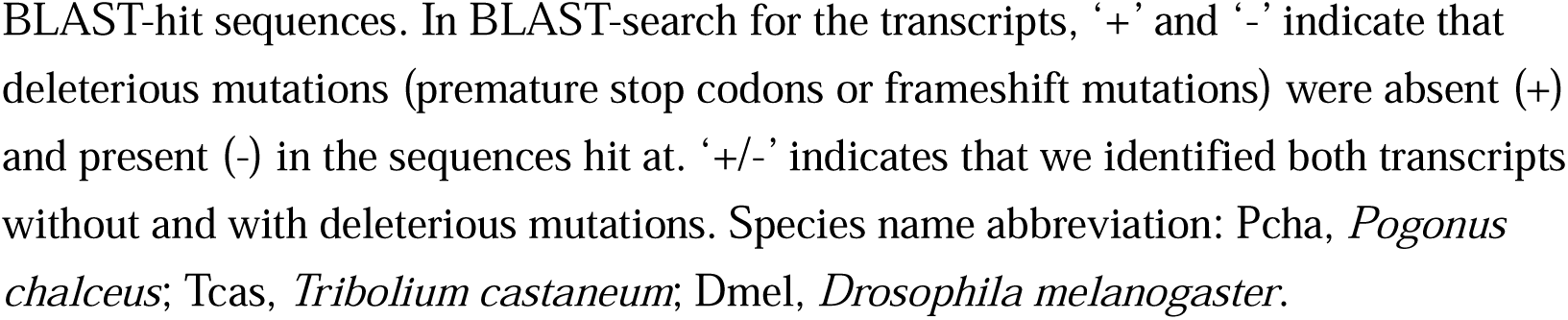
Photoreceptor and phototransduction genes detected in the assembled genome and transcripts.

| Query |  |  | *T. kuznetsovi* |  |
| --- | --- | --- | --- | --- |
| Protein name | Species | Accession no. | Genome | Transcripts |
| Visual opsin proteins |  |  |  |  |
| Lw opsin | Pcha | APY20654 | yes | yes (+) |
| Uv opsin | Pcha | APY20653 | yes | no |
| Non-visual opsin protein |  |  |  |  |
| C-opsin | Tcas | NP_001138950 | yes | yes (+) |
| Phototransduction proteins |  |  |  |  |
| Arr1 | Tcas | XP_966595 | yes | yes (+) |
| Arr2 | Tcas | NP_001164084 | yes | yes (+) |
| Gα49B | Tcas | XP_966311 | yes | yes (+/-) |
| Gβ76C | Tcas | XP_973851 | yes | yes (+/-) |
| Gγ30A | Tcas | XP_972123 | yes* | yes (+) |
| Gprk1 | Tcas | XP_966480 | yes | yes (+/-) |
| InaD | Tcas | XP_015837295 | yes | yes (+) |
| NinaA | Tcas | XP_973192 | yes | yes (+) |
| NinaC | Tcas | XP_015835531 | yes | yes (+) |
| NorpA | Tcas | XP_001812780 | yes | yes (+/-) |
| Pkc53E | Dmel | NP_476682 | yes | yes (+) |
| Rab6 | Tcas | XP_972453 | yes | yes (+) |
| RdgC | Tcas | XP_974915 | yes | yes (+/-) |
| Trp | Tcas | XP_968670 | yes | yes (+) |
| Trpl | Tcas | XP_968598 | yes | no |

'yes' indicates that the genes were detected at e-value $< 1 \times e^{-20}$ . 'yes\*' indicates that the gene was detected at e-value $\geq 1 \times e^{-20}$ . 'no' indicates that the genes were not detected in
BLAST-hit sequences. In BLAST-search for the transcripts, ‘+’ and ‘-’ indicate that deleterious mutations (premature stop codons or frameshift mutations) were absent (+) and present (-) in the sequences hit at. ‘+/-’ indicates that we identified both transcripts without and with deleterious mutations. Species name abbreviation: Pcha, *Pogonus chalceus*; Tcas, *Tribolium castaneum*; Dmel, *Drosophila melanogaster*.

### Opsin genes

One *lw opsin* gene of *T. kuznetsovi* was identified with BLAST search. The gene was divided into three genome scaffolds and two transcript contigs (Table S2). The scaffolds and contigs were joined together using PCR and RT-PCR, and then exon and intron regions of the *lw opsin* gene were determined (Fig. 2A). The conceptually translated Lw opsin amino acid sequence was 379 residues and consisted of six exons. There was neither a premature stop codon nor a frameshift mutation in the coding sequence.

**Fig. 2.**
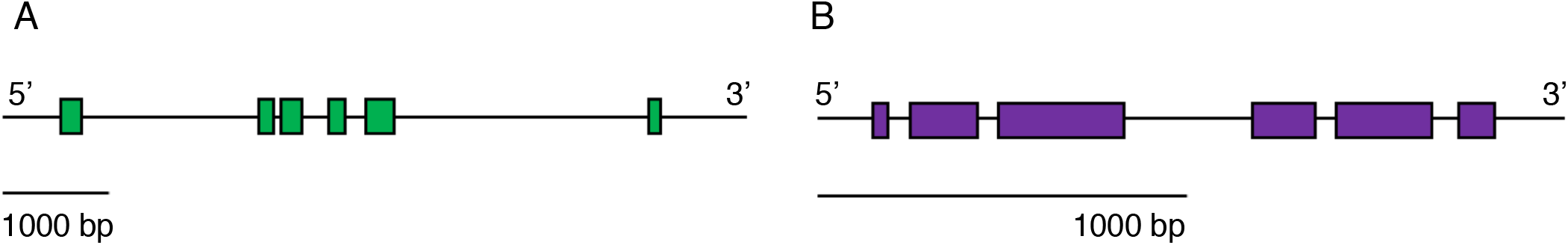
The structure of opsin genes in *T. kuznetsovi*. (A) The putative structure of *lw opsin* gene, which consists of six coding exons. (B) The putative structure *uv opsin* gene, which consists of six coding exons.

One *uv opsin* gene of *T*. *kuznetsovi* was identified by performing BLAST search. The gene was present within one scaffold in the genome and no contig was found in the transcripts (Table S2). The cDNA sequence of the *uv opsin* amino acid sequence was determined using RT-PCR, and then exon and intron regions of the *uv opsin* gene were determined (Fig. 2B). The conceptually translated Uv opsin amino acid sequence was 373 residues and consisted of six exons. There was neither a premature stop codon nor a frameshift mutation in the coding sequences.

### Opsin phylogeny and selective pressure

A molecular phylogenetic tree of opsin genes of *T. kuznetsovi, P. chalceus, G. marinus, T. marmoratus* and *T. castaneum* and *A. mellifera* was reconstructed (Fig. 3, Table S4). The *lw opsin* and *uv opsin* of *T. kuznetsovi* were clustered with those of *P. chalceus*, in accordance with their taxonomic relationship.

**Fig. 3.**
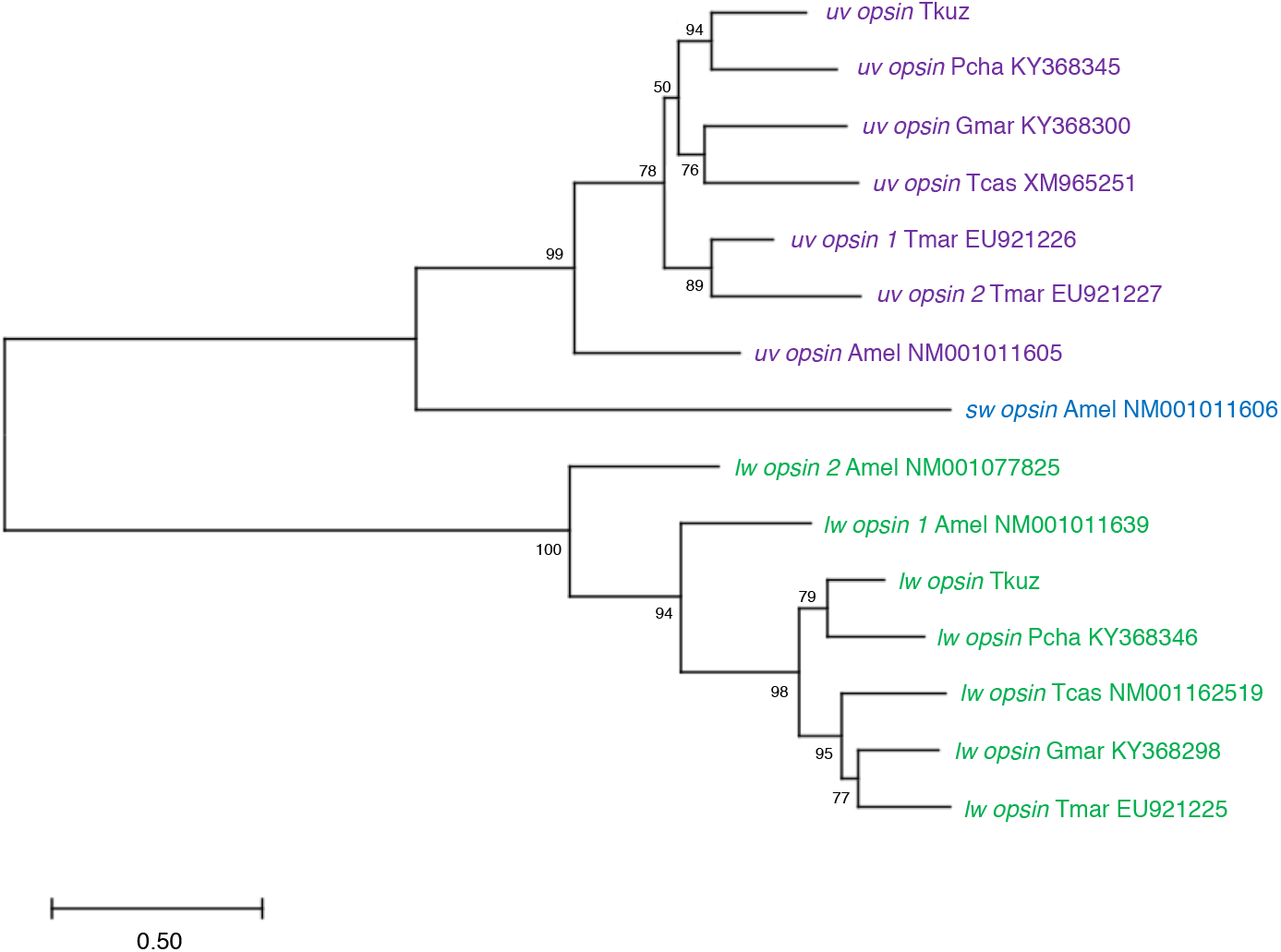
A maximum likelihood tree for visual opsin genes. Bootstrap probabilities are provided on nodes. OTU names consist of the abbreviated species names, gene names and accession numbers: Tkuz, *Trechiama kuznetsovi;* Pcha, *Pogonus chalceus;* Gmar, *Gyrinus marinus;* Tmar, *Thermonectus marmoratus;* Tcas, *Tribolium castaneum;* Amel, *Apis mellifera*.

In *lw opsin* genes, Ka/Ks ratio was 0.158131 between the sequences of the ancestor and *T. kuznetsovi* (*p* = 1.12862e^-013^, Fisher’s exact test), and Ka/Ks ratio was 0.0275072 between the sequences of the ancestor and *P*. *chalceus* (*p* = 2.70467e^-109^, Fisher’s exact test) (Table 3). In *uv opsin*, Ka/Ks ratio was 0.138044 between the sequences of the ancestor and *T*. *kuznetsovi* (*p* = 1.34285e^-024^, Fisher’s exact test), and Ka/Ks ratio was 0.0695552 between the sequences of the ancestor and *P*. *chalceus (p* = 7.58434e^-092^, Fisher’s exact test). In all of these cases, Ka/Ks ratios were far below 1.0, indicating that opsin genes have been under negative (purifying) selection in both the lineage leading to *T. kuznetsovi* and that leading to *P. chalceus*.

**Table 3.**
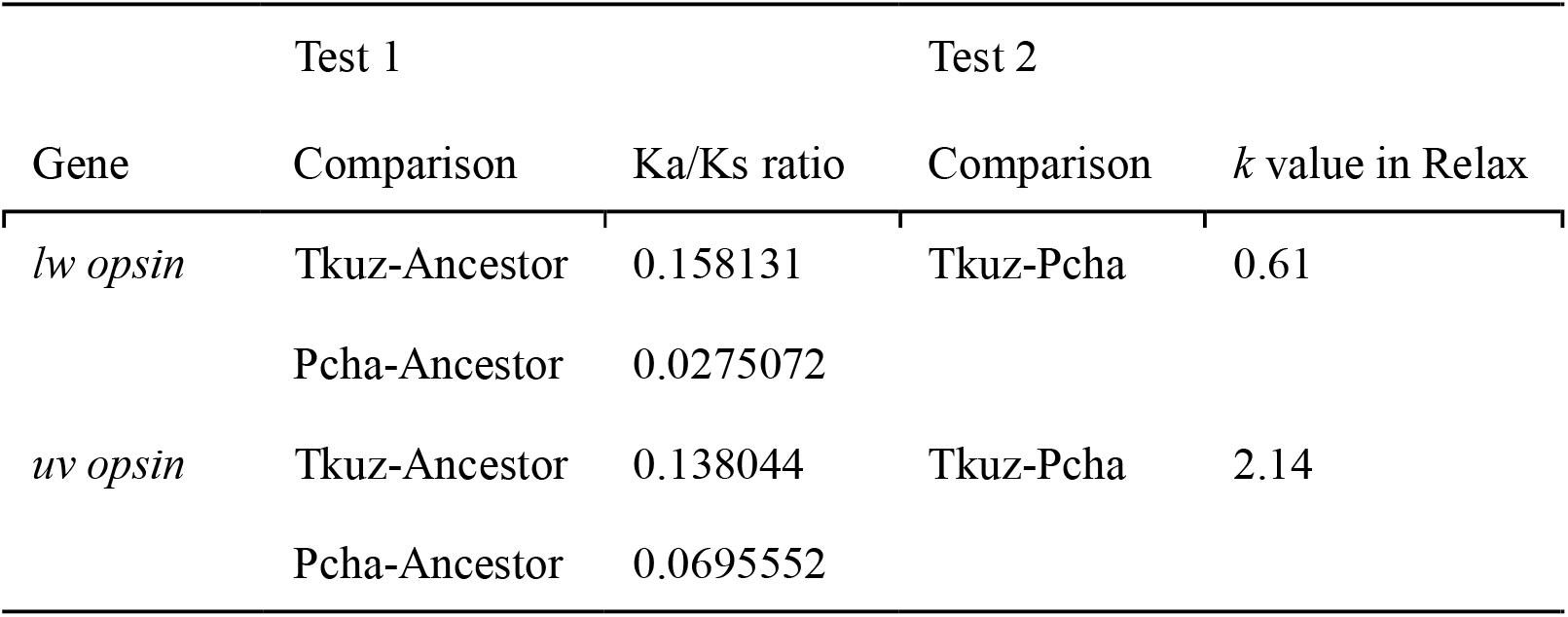
The test of selection in opsin genes of *T. kuznetsovi*.

Subsequently, the difference in the degree of the purifying selection between the lineages from the ancestor to *T. kuznetsovi* and to *P. chalceus* was tested. As the result of Relax analysis in Hyphy, *k* value was 0.61 (*p* = 0.225) in *lw opsin* and 2.14 (*p* = 0.539) in *uv opsin* (Table 3). Because this analysis did not show a statistical significance, we were unable to conclude whether the selection on opsin genes was relaxed or intensified in the lineage leading to *T. kuznetsovi* compared to the control lineage.

### Internal structure of a compound eye

Putative photoreceptor cells stained by hematoxylin were observed in the internal structure of a vestigial compound eye in a *T*. *kuznetsovi* adult (Fig. 4A). The surface was covered by a transparent cuticle, a cornea. By observation from the outer surface of the head, we could see a transparent cornea and ocular ridge with black pigmentation (Fig. 1D). There was no pigmentation in cells within the eye structure, unlike compound eyes of other carabid beetles [44]. No crystalline cones or any similar structure were found [45]. An optic stalk, which is a nerve bundle connecting a compound eye and a brain, was observed [46] (Fig. 4B).

**Fig. 4.**
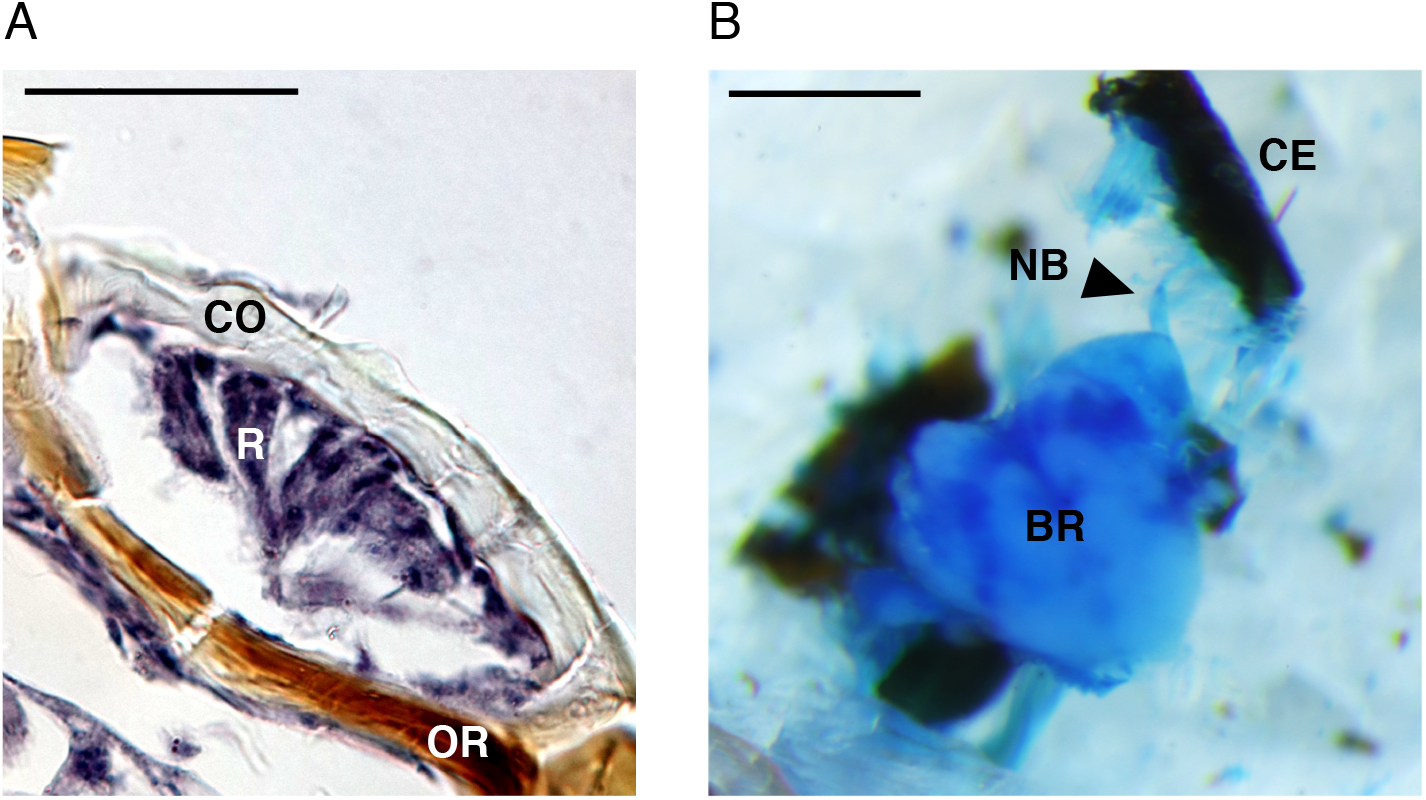
The internal structure and nerve tissue of *T. kuznetsovi* head. (A) Transverse section of a compound eye, dorsal is up. Scale bar indicates 20 μm. CO, cornea; R, putative photoreceptor cells; OR, ocular ridge. (B) Dorsal view of dissected head. Nerve tissue was stained with methylene blue solution. Scale bar indicates 0.5 mm. CE, compound eye; NB, nerve bundle; BR, brain (supraesophageal ganglion).

## Discussion

To understand the process of subterranean colonization of organisms, the question of whether shallow subterranean habitats are a gateway to colonizing deep zones has been featured in subterranean biology [3, 4]. In previous studies, organisms inhabiting a cave or a calcrete aquifer and having extremely regressed structure and function of vision were examined [8, 11–14]. To understand the transitional stage in subterranean evolution, we focused on a trechine beetle, *Trechima kuznetsovi*, which inhabits the upper hypogean zone and has a vestigial compound eye [16]. We evaluated the ability of photoreception in *T. kuznetsovi* by genomics and histological observation.

We identified one *lw opsin* gene and one *uv opsin* gene in the genome and in the transcripts in the adult. No frameshift mutation or premature stop codon was found in these exon regions, Ka/Ks ratios were less than 1.0, and there was no significant difference in the selective pressure between evolutionary lineages of subterranean *T. kuznetsovi* and surface *P. chalceus*. These analyses implied that Lw opsin and Uv opsin are under functional constraint, which could be one piece of evidence supporting the possibility that *T. kuznetsovi* has photoreception ability for long wavelength and ultraviolet light. Transcripts of 15 phototransduction genes without deleterious mutations (premature stop codons or frameshift mutations) were detected in the assembled transcripts. One phototransduction gene, *trpl*, was found in the assembled genome. These results suggested the ability of phototransduction of *T. kuznetsovi*.

In subterranean diving beetles in Western Australia, the *lw opsin* gene became a pseudogene due to frameshift mutations, and neither *lw opsin* nor *uv opsin* transcripts were observed [12–14]. Frameshift mutations and premature stop codons occurred in some phototransduction genes: *Arr1, Arr2, ninaC, trp* and *trpl* [14]. There are two possible causes for these differences between *T. kuznetsovi* and the subterranean diving beetles. The first possibility is the difference in their ecological niches. The calcrete aquifer, in which subterranean diving beetles live, is located at a depth of 10–30 m underground [11, 47]. In contrast, the upper hypogean zone, in which *T. kuznetsovi* adults live, is a few or some dozen centimeters below the slope surface of a v-shaped valley. *Trechiama kuznetsovi* adults would occasionally be exposed to the surface due to landslides occurring as a result of precipitation or earthquakes [48–50]. This temporary light stimulus could function as a selective pressure to retain the function of opsin genes and other phototransduction genes. Friedrich et al. [29] also suggested that the retention of photoreceptor genes and phototransduction genes in *Ptomaphagus hirtus* (Leiodidae) inhabiting the Frozen Niagara cave entrance of Mammoth Cave in Kentucky, USA. Beetles in shallow subterranean environments (the upper hypogean zone and cave entrance) could tend to maintain the photoreception. The second possibility is that opsin genes and other phototransduction genes of *T. kuznetsovi* have not spent enough time to become pseudogenes. Niemiller et al. [51] showed that some cave lineages in Amblyopsidae still possess functional rhodopsin, although they inhabit an aphotic environment. This retained functionality is thought to be due to insufficient accumulation of mutations during recent subterranean colonization. As in these cave lineages, pseudogenization of opsin genes in *T. kuznetsovi* could not be observed because divergence between *T. kuznetsovi* and related terrestrial species occurred recently. To further examine this possibility, the divergence time of *Trechiama* species needs to be studied.

By performing paraffin sectioning and dissection, we observed the cells inside the compound eye and the optic stalk connecting the compound eye and the brain in *T. kuznetsovi*. These observations suggested that photoreception is structurally possible even with the vestigial compound eyes of this species. Complete loss of compound eyes and optic lobes was observed in *Sinaphaenops wangorum* (Trechinae) inhabiting the deep area of a cave in Guangxi Autonomous Region, China [52]. Thus, the existence of the optic stalk is thought to be due to the retention of photoreception ability by *T. kuznetsovi*, not to an inability of the visual system to degenerate any further.

Collectively, the results of genomics and histology performed here suggested the ability of photoreception in *T. kuznetsovi*. This species is thought to possess both a surface trait (photoreception) and some subterranean traits (vestigial compound eye, underdeveloped body pigmentation and other morphological adaptations) [53, 54]. These characteristics would reflect an intermediate phase toward colonizing a deeper subterranean niche. Further understanding of the visual degeneration process will be obtained by clarifying the phylogenetic relationship between subterranean species and surface species of trechine beetles.

## Conclusions

By *de novo* assembly of genome and transcript sequences, we identified photoreceptor genes and phototransduction genes of a trechine beetle, *Trechiama kuznetsovi*, which inhabits the upper hypogean zone. The encoded amino acid sequences of *lw opsin* and *uv opsin* had neither a premature stop codon nor a frameshift mutation, and appeared to be subject to purifying selection. We identified potential photoreceptor cells in the compound eye and nerve bundle connected to the brain. The present findings suggest that *T. kuznetsovi* has retained the ability of photoreception.

## Abbreviations

Lw opsin: Long wavelength opsin
Uv opsin: Ultraviolet opsin
Arr1: Arrestin 1
Arr2: Arrestin 2
G(/49B: G protein α-subunit 49B
Gβ76C: G protein β-subunit 76C
Gy30A: G protein γ-subunit 30A
Gprk1: G protein-coupled receptor kinase 1
InaD: Inactivation no afterpotential D
NinaA: Neither inactivation nor afterpotential A
NinaC: Neither inactivation nor afterpotential C
NorpA: No receptor potential A
Pkc53E: Protein C kinase 53E
Rab6: Rab-protein 6
RdgC: Retinal degeneration C
Trp: Transient receptor potential
Trpl: Trp-like
CO: Cornea
R: Putative photoreceptor cells
OR: Ocular ridge
CE: Compound eye
NB: Nerve bundle
BR: Brain

## Declarations

### Ethical approval and consent to participate

Not applicable.

### Consent for publication

Not applicable.

### Availability of data and material

Sequence reads and scaffolds of the genome and transcripts of *Trechiama kuznetsovi* were deposited to DDBJ/EMBL/GenBank (DRR415324, DRR415325).

### Authors contributions

Conceptualization, T.N. and S.K.; Data acquisition, T.N., Y.T., and H.A.; Data analysis, T.N., Y.T, H.A. and S.K.; Writing, T.N. and S.K.; All authors approved the final manuscript.

## Acknowledgements

We thank Takashi Hayakawa and Juno Shimada for technical help; Cédric Finet for technical advice; Masahiro Ôhara for collection site information; Elizabeth Nakajima for English editing; and members of the Koshikawa lab for sample collection and discussion. The super-computing resource was provided by Human Genome Center (the Univ. of Tokyo).

---

Table S1.

The result of genome and transcript sequencing in *T. kuznetsovi*.

Table S2.

BLAST search for opsin genes to the assembled genome and transcripts in *T. kuznetsovi*.

Table S3.

Primers used in PCR and RT-PCR to amplify opsin genes in *T. kuznetsovi. T* (□), annealing temperature.

Table S4.

BLAST search for opsin genes to non-redundant protein sequences in NCBI database.

