## Supplementary material for "Photoreceptor genes in a trechine beetle, *Trechiama kuznetsovi*, living in the upper hypogean zone": Supplmental Tables

### Slide 1
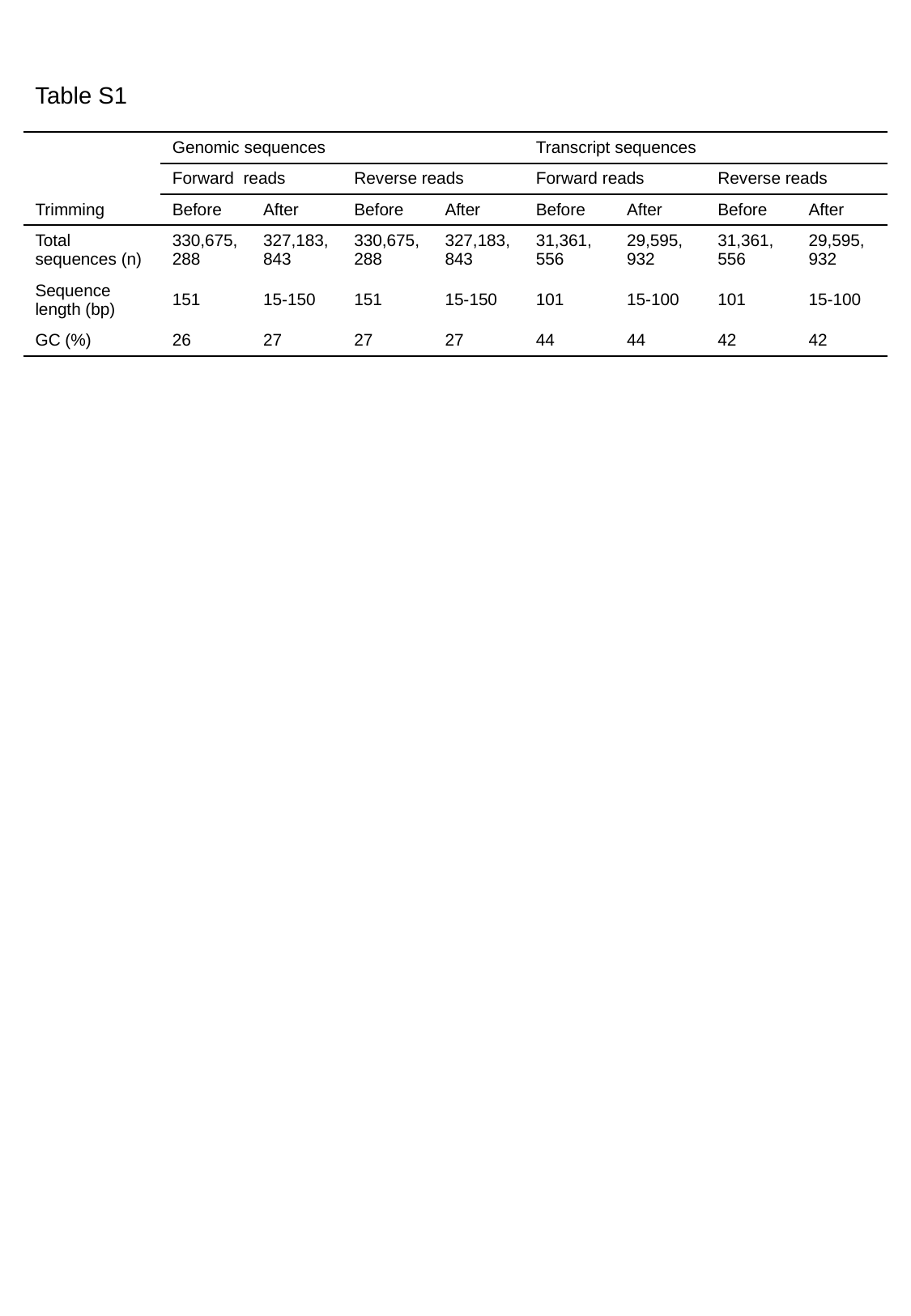

Table S1
| Trimming | Genomic sequences | | | | Transcript sequences | | | |
| --- | --- | --- | --- | --- | --- | --- | --- | --- |
| | Forward reads | | Reverse reads | | Forward reads | | Reverse reads | |
| Trimming | Before | After | Before | After | Before | After | Before | After |
| Total sequences (n) | 330,675, 288 | 327,183, 843 | 330,675, 288 | 327,183, 843 | 31,361, 556 | 29,595, 932 | 31,361, 556 | 29,595, 932 |
| Sequence length (bp) | 151 | 15-150 | 151 | 15-150 | 101 | 15-100 | 101 | 15-100 |
| GC (%) | 26 | 27 | 27 | 27 | 44 | 44 | 42 | 42 |

### Slide 2
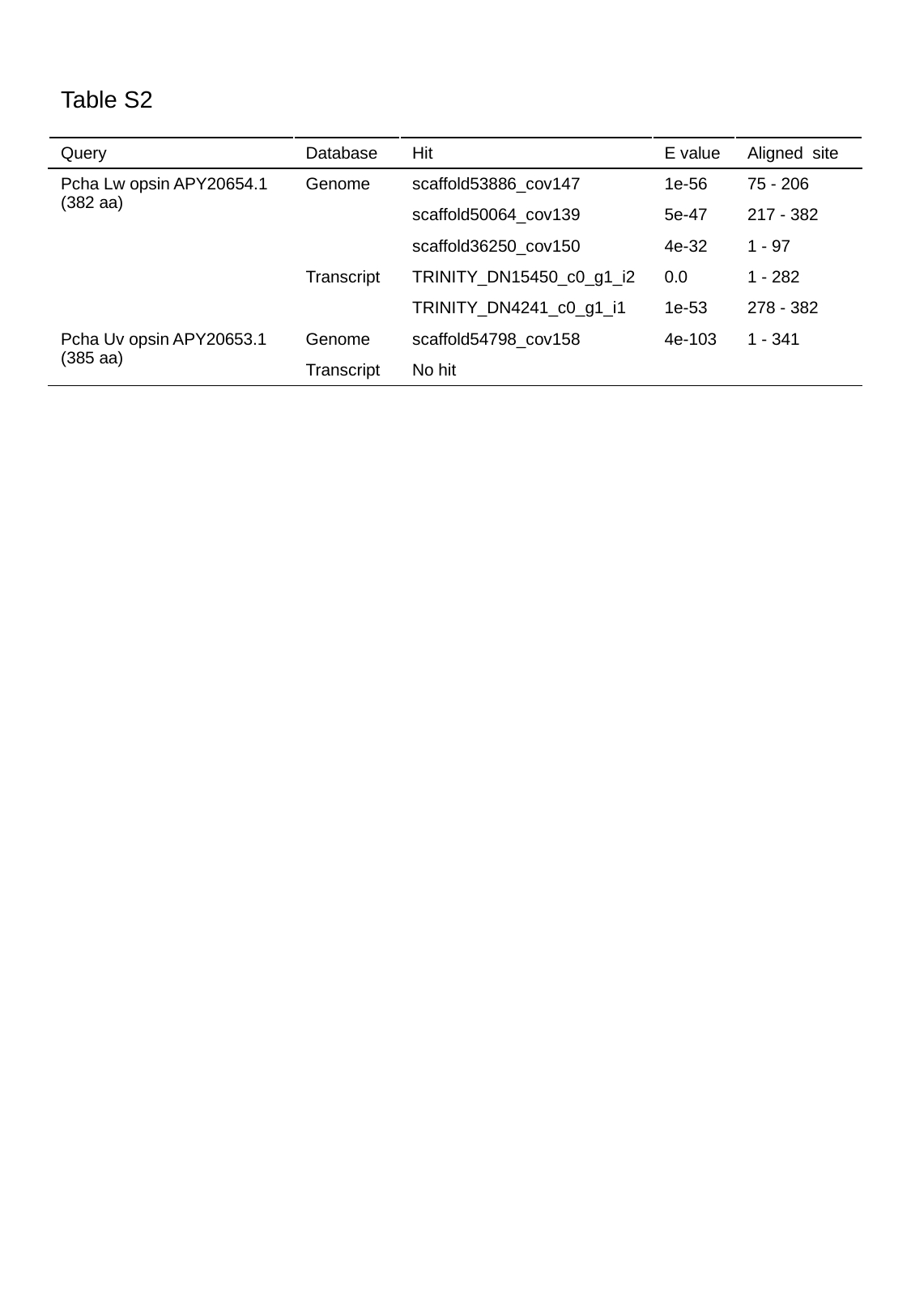

Table S2
| Query | Database | Hit | E value | Aligned site |
| --- | --- | --- | --- | --- |
| Pcha Lw opsin APY20654.1 (382 aa) | Genome | scaffold53886\_cov147 | 1e-56 | 75 - 206 |
| | | scaffold50064\_cov139 | 5e-47 | 217 - 382 |
| | | scaffold36250\_cov150 | 4e-32 | 1 - 97 |
| | Transcript | TRINITY\_DN15450\_c0\_g1\_i2 | 0.0 | 1 - 282 |
| | | TRINITY\_DN4241\_c0\_g1\_i1 | 1e-53 | 278 - 382 |
| Pcha Uv opsin APY20653.1 (385 aa) | Genome | scaffold54798\_cov158 | 4e-103 | 1 - 341 |
| | Transcript | No hit | | |

### Slide 3
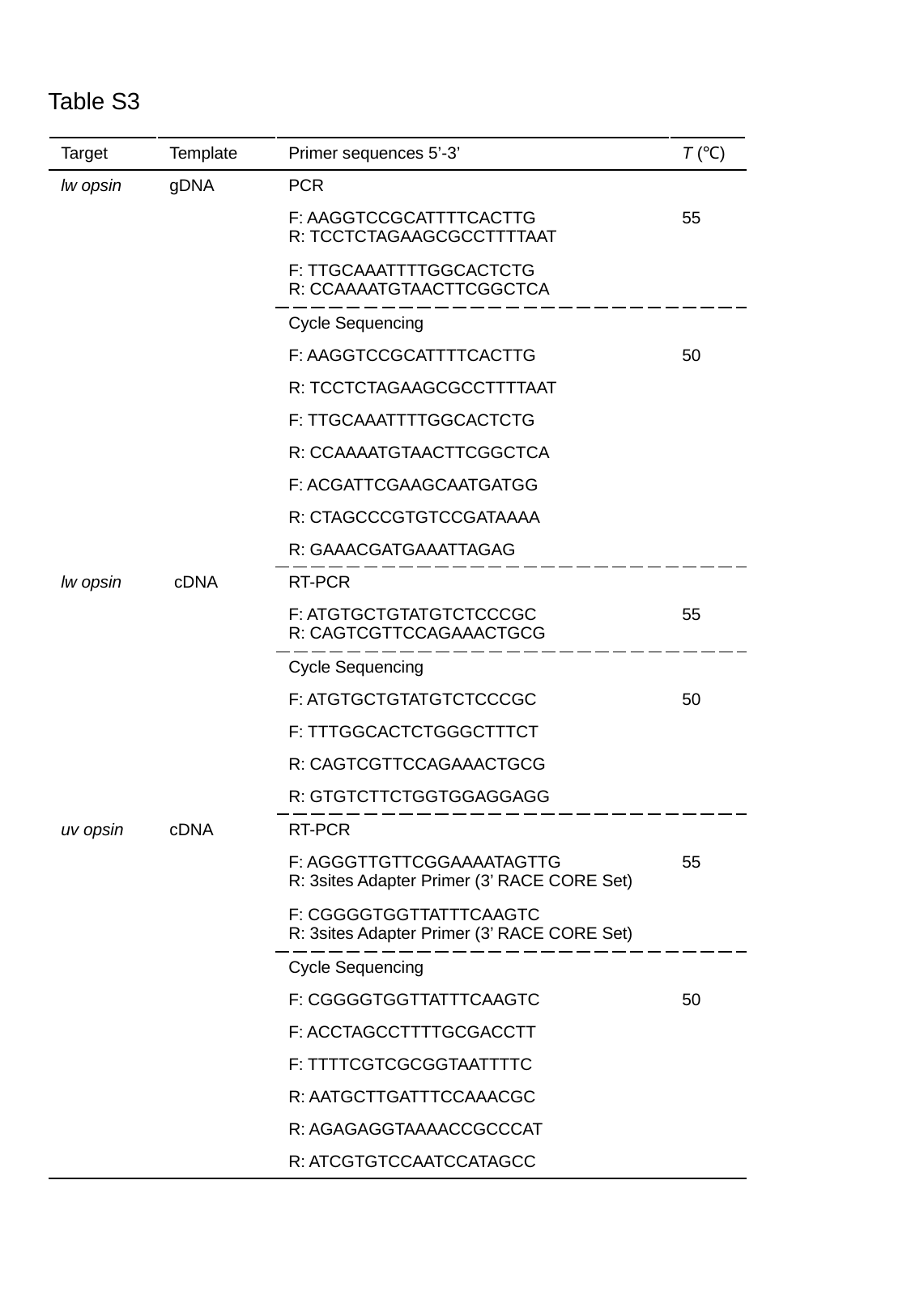

Table S3
| Target | Template | Primer sequences 5’-3’ | T (℃) |
| --- | --- | --- | --- |
| lw opsin | gDNA | PCR | |
| lw opsin | gDNA | F: AAGGTCCGCATTTTCACTTG R: TCCTCTAGAAGCGCCTTTTAAT | 55 |
| | | F: TTGCAAATTTTGGCACTCTG R: CCAAAATGTAACTTCGGCTCA | |
| | | Cycle Sequencing | |
| | | F: AAGGTCCGCATTTTCACTTG | 50 |
| | | R: TCCTCTAGAAGCGCCTTTTAAT | |
| | | F: TTGCAAATTTTGGCACTCTG | |
| | | R: CCAAAATGTAACTTCGGCTCA | |
| | | F: ACGATTCGAAGCAATGATGG | |
| | | R: CTAGCCCGTGTCCGATAAAA | |
| | | R: GAAACGATGAAATTAGAG | |
| lw opsin | cDNA | RT-PCR | |
| lw opsin | cDNA | F: ATGTGCTGTATGTCTCCCGC R: CAGTCGTTCCAGAAACTGCG | 55 |
| | | Cycle Sequencing | |
| | | F: ATGTGCTGTATGTCTCCCGC | 50 |
| | | F: TTTGGCACTCTGGGCTTTCT | |
| | | R: CAGTCGTTCCAGAAACTGCG | |
| | | R: GTGTCTTCTGGTGGAGGAGG | |
| uv opsin | cDNA | RT-PCR | |
| uv opsin | cDNA | F: AGGGTTGTTCGGAAAATAGTTG R: 3sites Adapter Primer (3’ RACE CORE Set) | 55 |
| | | F: CGGGGTGGTTATTTCAAGTC R: 3sites Adapter Primer (3’ RACE CORE Set) | |
| | | Cycle Sequencing | |
| | | F: CGGGGTGGTTATTTCAAGTC | 50 |
| | | F: ACCTAGCCTTTTGCGACCTT | |
| | | F: TTTTCGTCGCGGTAATTTTC | |
| | | R: AATGCTTGATTTCCAAACGC | |
| | | R: AGAGAGGTAAAACCGCCCAT | |
| | | R: ATCGTGTCCAATCCATAGCC | |

### Slide 4
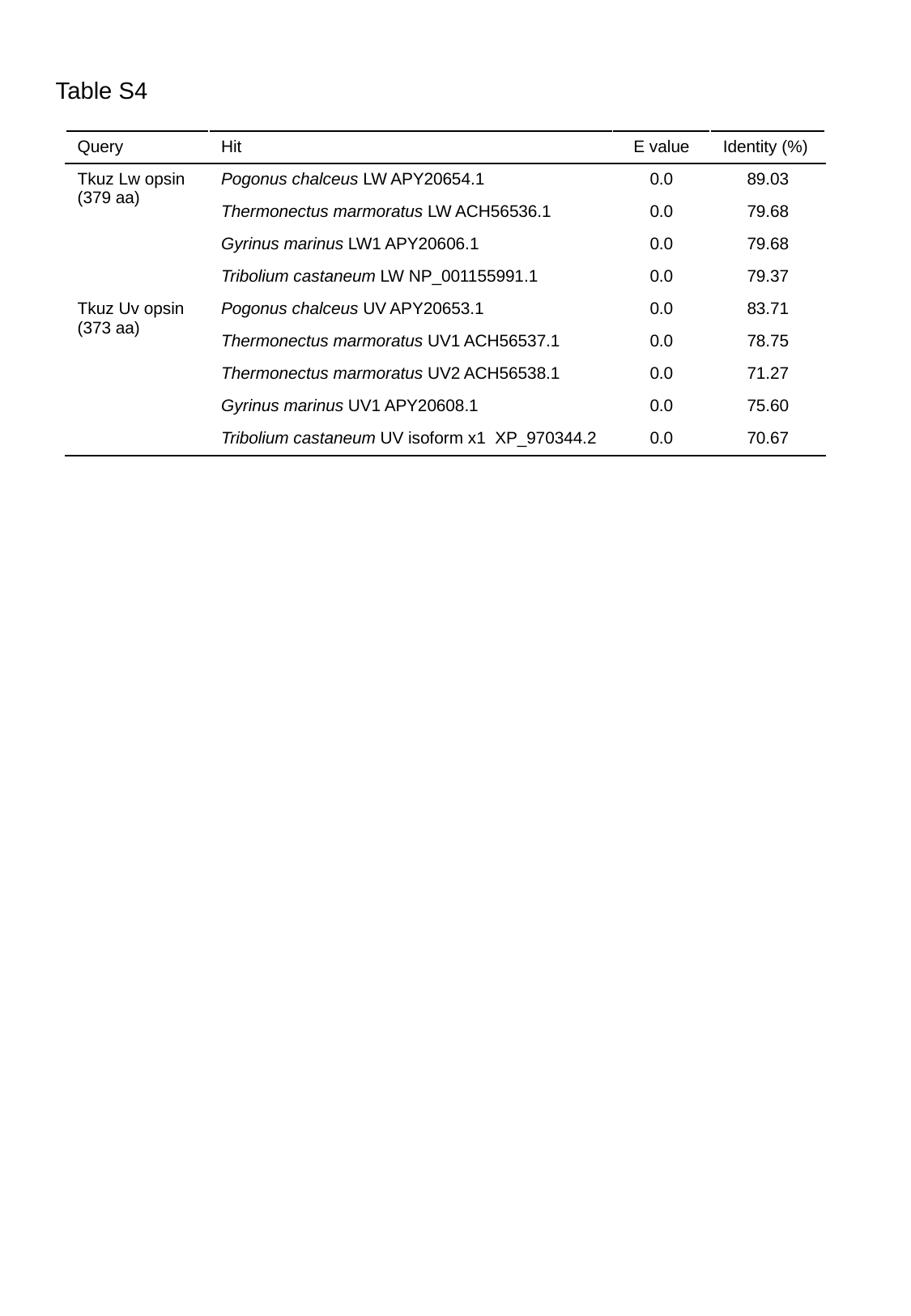

Table S4
| Query | Hit | E value | Identity (%) |
| --- | --- | --- | --- |
| Tkuz Lw opsin (379 aa) | Pogonus chalceus LW APY20654.1 | 0.0 | 89.03 |
| | Thermonectus marmoratus LW ACH56536.1 | 0.0 | 79.68 |
| | Gyrinus marinus LW1 APY20606.1 | 0.0 | 79.68 |
| | Tribolium castaneum LW NP\_001155991.1 | 0.0 | 79.37 |
| Tkuz Uv opsin (373 aa) | Pogonus chalceus UV APY20653.1 | 0.0 | 83.71 |
| | Thermonectus marmoratus UV1 ACH56537.1 | 0.0 | 78.75 |
| | Thermonectus marmoratus UV2 ACH56538.1 | 0.0 | 71.27 |
| | Gyrinus marinus UV1 APY20608.1 | 0.0 | 75.60 |
| | Tribolium castaneum UV isoform x1 XP\_970344.2 | 0.0 | 70.67 |
